# An analysis of contractile and protrusive cell behaviours at the superficial surface of the zebrafish neural plate

**DOI:** 10.1101/2024.02.20.581322

**Authors:** Claudio Araya, Raegan Boekemeyer, Francesca Farlie, Lauren Moon, Freshta Darwish, Chris Rookyard, Leanne Allison, Gema Vizcay-Barrena, Roland Fleck, Millaray Aranda, Masa Tada, Jonathan D W Clarke

**Affiliations:** Laboratory of Developmental Biology, Instituto de Ciencias Marinas y Limnológicas, Facultad de Ciencias, Universidad Austral de Chile, Campus Isla Teja, 5090000, Valdivia, Chile; MRC Centre for Developmental Neurobiology, King’s College London, New Hunt’s House, 4th Floor, Guy’s Hospital Campus, London, United Kingdom; Centre for Ultrastructural Imaging, King’s College London, New Hunt’s House, 4th Floor, Guy’s Hospital Campus, London, United Kingdom; Department of Cell and Developmental Biology, University College London, WC1E 6BT London, United Kingdom

## Abstract

The motive forces underlying convergence and internalisation of the teleost neural plate remain unknown. To help understand this crucial morphogenetic movement we have analysed collective and individual cell behaviours at the superficial surface of the neural plate. Convergence to the midline does not rely on mediolateral cell intercalation, it is characterised by oscillatory contractile behaviours, a punctate distribution of Cdh2 and medially polarised actin-rich protrusions at the surface of the neural plate. Additionally, we characterise and discuss the intimate relationship and dynamic cell surfaces between the motile neural plate and the stationary overlying non-neural enveloping layer.

## Introduction

The early stages of neural tube formation are characterized by collective cell activities effecting neural plate narrowing and internalisation beneath the protective non-neural ectoderm. In the past years, accumulating evidence shows that convergent and extension movements and apical constriction are the two dominant cell behaviors responsible for this neural plate shaping (Colas and Schoenwolf, 2001; Nikolopoulou et al., 2017). Convergent and extension through mediolateral cell intercalation facilitates neural plate narrowing (Wallingford and Harland, 2002; Nishimura et al., 2012; Williams et al., 2014; Butler and Wallingford, 2018), while apical cell constriction is a key behavior driving tissue internalisation around the dorsal midline (Haigo et al., 2003; Nishimura and Takeichi, 2008). In addition, tensile forces generated within and around the neuroepithelium appears as key regulator for neural tube morphogenesis (Galea et al., 2017; 2021; Marshall et al., 2023). The precise cellular and sub-cellular details of these neural plate cell dynamics remain poorly understood.

In recent years, live imaging studies in invertebrate systems have revealed that collective cell deformation is not driven by smooth continuous cell shape changes but through oscillatory steps of cell contractions (Solon et al., 2009; Kim and Davidson, 2011; Martin and Goldstein, 2014; Munjal et al., 2015; Gorfinkiel, 2016; Coravos et al., 2017; Sun and Toyoma, 2018). Key to this oscillatory cell behavior is a dynamic actomyosin network, which confers pulsatile tensile forces during cell and tissue remodeling (Martin et al., 2009; 2010; Rauzi et al., 2010; Gorfinkiel and Blanchard, 2011; Mason et al., 2013; Munjal et al., 2015; An et al., 2017). Furthermore, junctional cell-cell adhesion components like classical Cadherins appear as critical components integrating actomyosin contractile forces at both cell and tissue level (Martin et al., 2010). Thus, coupled pulsatile contractions have emerged as a key driver for invertebrate tissue remodeling, and although less deeply understood during vertebrate morphogenesis, oscillatory cell contractility appears to be a conserved feature in vertebrate tissue remodelling (Kim and Davidson, 2011; Nicolas-Perez et al., 2016; Werner et al., 2021; Christodoloulou and Skourides, 2022).

In this study we aim to improve our understanding of vertebrate tissue narrowing and internalisation by characterizing these movements at cell and subcellular levels using *in vivo* imaging in the zebrafish neural plate. The zebrafish neural primordium narrows dramatically and internalises? during its transition from neural plate to neural keel, and although the teleost neural plate has a different cytoarchitecture compared to other vertebrates (reviewed in Araya et al., 2016) it uses several morphogenetic mechanisms conserved with other vertebrates (Ciruna et al., 2006; Hong and Brewster, 2006; Tawk et al., 2007; Araya et al., 2019; Araya and Carrasco, 2021; Werner et al., 2021).

In common with amniote and other anamniote embryos zebrafish neural plate convergence depends on the activity of the non-canonical Wnt/Planar Cell Polarity (PCP) signaling pathway (Copp et al., 2003; Ciruna et al., 2006; Wallingford, 2006; Wang et al., 2006; Tawk et al., 2007; Nishimura et al., 2012; Williams et al., 2014; Butler and Wallingford, 2018;). The PCP pathway cannonically drives convergence through mediolateral cell intercalation (Ybot-Gonzalez et al., 2007; Williams et al., 2014), but this hasn’t been shown in the fish neural plate. In addition, zebrafish neural plate convergence and internalisation is critically dependent on the function of the cell-cell adhesion protein Cdh2 (Lele et al., 2002; Hong and Brewster, 2006; Biswas et al., 2010; Araya et al., 2019). At the cellular level, zebrafish neural plate cells adopt cell surface constriction strategies typical of vertebrate epithelia in order to effectively reduce their superficial surface and internalise. Myosin-II is a critical component for superficial cell surface deformation which also depends on Cdh2 activity (Araya et al., 2019). Abrogation of Cdh2 results in defective Myosin-II distribution, mislocalised cell internalisation events and defective neural plate morphogenesis (Araya et al., 2019).

In this study, we used high spatial and rapid temporal *in vivo* imaging to study cell surface dynamics during zebrafish neural plate convergence and internalisation in prospective hindbrain regions. We find that during the neural plate to neural keel transition, neural plate narrows as a coherent sheet of cells without conventional mediolateral intercalation events. Analysis reveals that convergence is characterised by medially orientated actin-rich protrusions that lie on the surface of the neural plate and a punctate distribution of Cdh2 between superficial cell interfaces. Both convergence and internalisation are accompanied by oscillatory contractile behaviours at superficial neural plate cell surfaces. Cdh2 is also required for the medially polarised protrusive activity of neural plate cells. Finally, we analyse the interface and cell protrusions between neural plate and the overlying non-neural enveloping layer (the EVL), that suggest the EVL might play a role in neural plate morphogenesis.

## Results

### Cell rearrangements during convergence to midline

To understand the movements of the neural plate during convergence and internalisation we have used time-lapse confocal imaging to monitor the relative movements of cells at the neural plate surface in the region of the developing hindbrain between 10 and 12 hpf (hours post fertilisation). Images were acquired from the dorsal surface of the neural plate. We analysed cohorts of approximately 50 cells for neighbour exchange and asked whether groups of cells exhibited any of the canonical characteristics of convergent extension as they converged towards the midline. We found cohorts of adjacent cells exhibited a small amount of mediolateral compression (approximately 20%) and a small amount of anteroposterior compression (approximately 10%, Fig. 1A-C, Movie 1). The overall area of the cohort of cells decreased by approximately 30% (Figure 1d). Cells move with a fairly constant speed of approximately 1 µm.min. Moreover individual cells mostly maintained neighbour relations during this movement and mediolateral intercalation rearrangements were very rare (Fig. 1A,B,F,G).

**Figure. 1.**
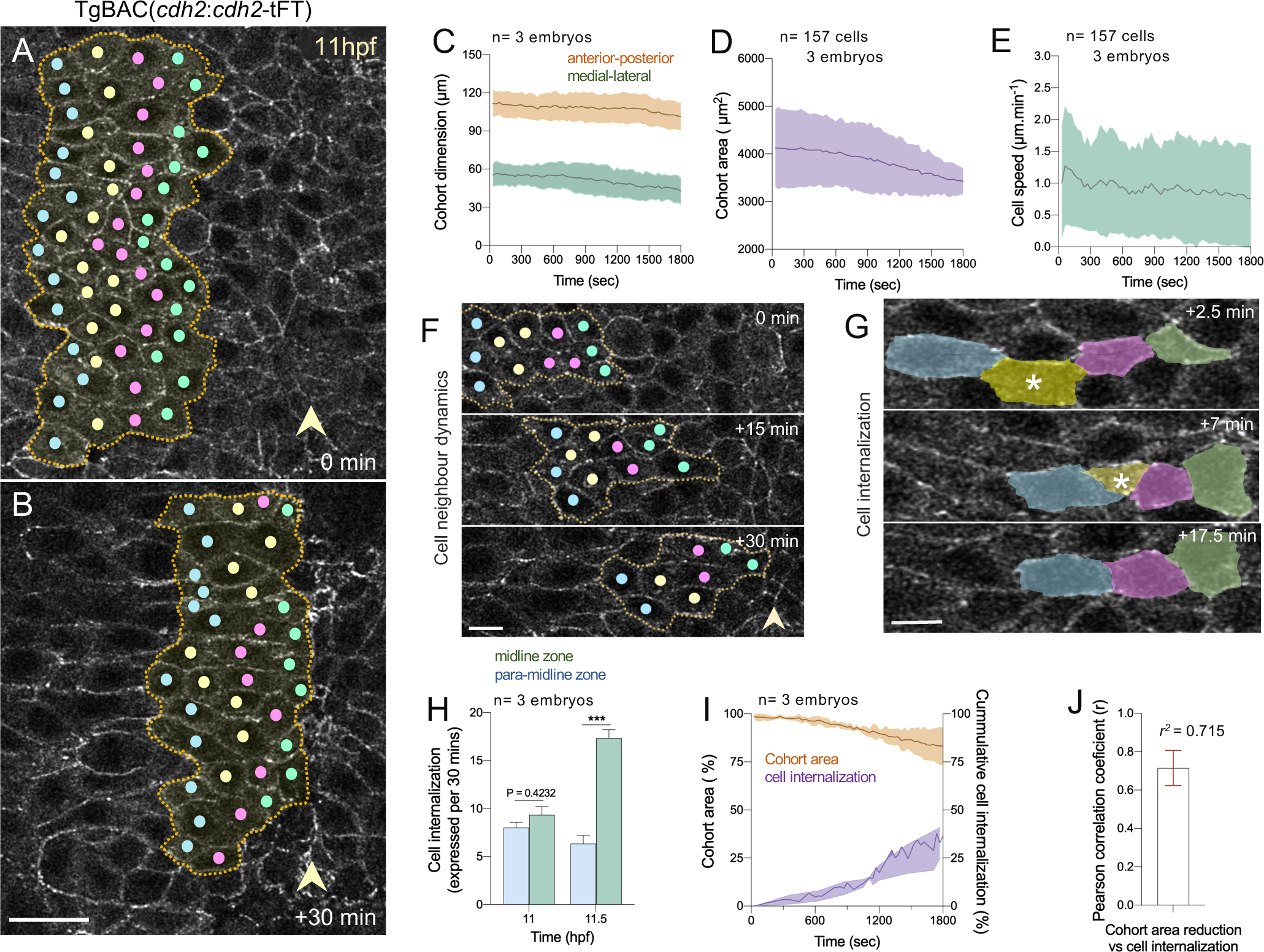
Neural plate cell dynamics during zebrafish midline convergence. (A, B) Representative *en face* view of single confocal stills from a time-lapse recording from a Cdh2-GFP transgenic embryo between 11-11.5hpf. Coloured dots indicate the centroids of individual neural plate cells as they move towards the dorsal midline. Dotted lines denote the outline of a cohort of cells. Arrowheads highlight the position of neural plate midline. Up is anterior and right is medial. Scale bar 20µm. (C) Cohort dimension changes during convergence (mean ± s.d., 3 embryos, average cohort n=52 cells). (D) Change in area of 3 cohorts of cells during midline convergence (mean ± s.d.). (E) Cell speed during convergence (mean ± s.d.). (F) Example of cell neighbour relations during convergence. Scale bar 10µm. (G) Example of cell internalisation event (asterisk) during convergence. Scale bar 10µm. (H) Cell internalisation events in midline zone (pale blue) and the para-midline zone (green) during convergence (****P<0.0001 Student’s t-test; mean ± s.e.m.). (I) Cohort area change (µm^2^) vs cell internalisation events during midline convergence. Pearson correlation r^2^ = 0.715±0.158; mean ± s.d.; n=3 embryos.

Once cells arrive to within a few cell’s widths of the midline, they no longer move as a cohort, rather cells individually internalise in a scattered fashion (Fig. 1F,G) as previously reported (Araya et al., 2019). The midline can be recognised by a high concentration of cells with small surface profiles. Approximately 30% of cells internalise in the period of our analysis (Fig. 1H,I), and there is a significant correlation between reduced cohort area and internalisation (Fig. 1J).

### Distribution and dynamics of Cdh2 and actin rich protrusions at the superficial NP surface

Our previous work has shown a relative accumulation of the adhesion molecule Cdh2 and the actomyosin motor complex at the superficial surface of neural plate cells. Furthermore both Cdh2 and actomyosin activity are critical requirements for neural plate movements (Araya et al., 2019). To better understand their contribution to tissue and cell movements we have used live reporters and high resolution *in vivo* imaging of their distribution and dynamics.

Imaging of TgBAC(*cdh2*:*cdh2*-tFT) embryos shows the superficial surface of neural plate cells is characterised by a punctate distribution of Cdh2 clusters. Membranes express a lower level of Cdh2 between the puncta. The puncta evenly encircle the superficial surface of individual cells, sitting at the interface with neighbouring cells (Fig. 2A-D). By imaging thin volumes of the superficial neural plate surface with a frame interval of between 5 and 10 seconds, we find the puncta are quite dynamic, appearing and disappearing, and sometimes apparently fusing with neighbouring puncta (Fig. 2C and Movies 2 and 3). Some puncta appear to be present on small motile protrusions that sit at the periphery of the cell surfaces (Fig. 2D). Cdh2-GFP puncta are more prominent at the superficial surface of neural plate cells than deeper surfaces of the cell where the fluorescence signal is lower and more evenly spread around the cell membranes (Fig. 2E).

**Figure. 2.**
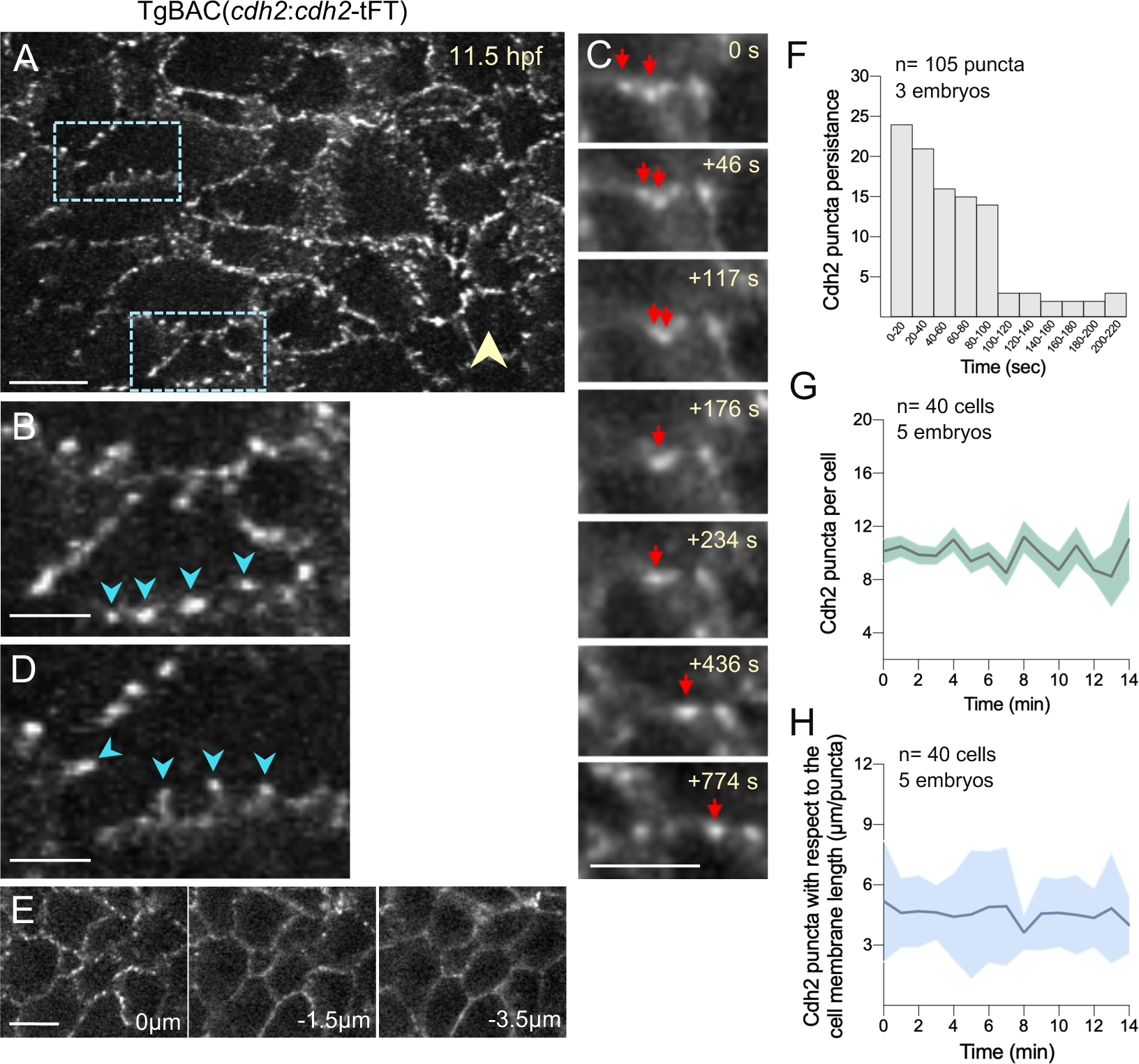
Cdh2 sub-cellular dynamics during midline convergence. (A) *En face* view of dorsal surface of neural plate cells (maximum projection of three z-levels) from a Cdh2-GFP embryo at 11.5 hpf. Yellow arrowhead indicates the midline. Anterior is up, right is medial. Scale bar 10µm. (B) Zoom of inset from a, depicting punctate organization of Cdh2-GFP (blue arrowheads). Scale bar 5µm. (C) Stills taken from a time-lapse of Cdh2-GFP embryos showing apparent fusion event between adjacent puncta (red arrows). Scale bar 10µm. (D) Zoom of inset from A, depicting Cdh2-GFP rich puncta on projections from cell membrane (blue arrowheads). Scale bar 5µm. (E) Three z-levels from surface of Cdh2-GFP neural plate. (F) Persistence of Cdh2-puncta at neural plate surface (n=105 puncta analysed from 3 embryos). (G) Quantification of absolute number of Cdh2-GFP puncta at the cell boundary during midline convergence. The average number of Cdh2-puncta present at the cell membrane during midline convergence is 9.834± 0.954 (n = 40 cells analysed from 3 embryos; mean ± s.d).

By imaging the actin binding protein utrophin using the Tg(*actb1:GFP-utrCH*) transgenic line, we find the superficial surface of the neural plate is covered in extremely dynamic actin-rich filopodia and lamellipodial protrusions. When the whole population of cells is imaged these protrusions are seen around the superficial perimeter of every cell (Movie 4). However when the tissue is mosaically labelled for utr-RFP it is clear the actin rich protrusions have a very distinct polarity as they predominantly protrude medially towards the midline (Fig. 3A, B). The protrusions lie over the surface of the neighbouring (usually more medial) cell and move dynamically on this surface (Movie 5). Medially directed protrusions are confirmed by serial block face imaging and this further shows these protrusions lie in the plane of the interface between neural plate and the overlying enveloping layer (EVL) (Fig. 3D and see later Fig. 8H, I). By imaging Cdh2 and utr in the same cells we find that the actin rich protrusions extend beyond than the Cdh2 puncta (Fig. 3E, F).

**Figure 3.**
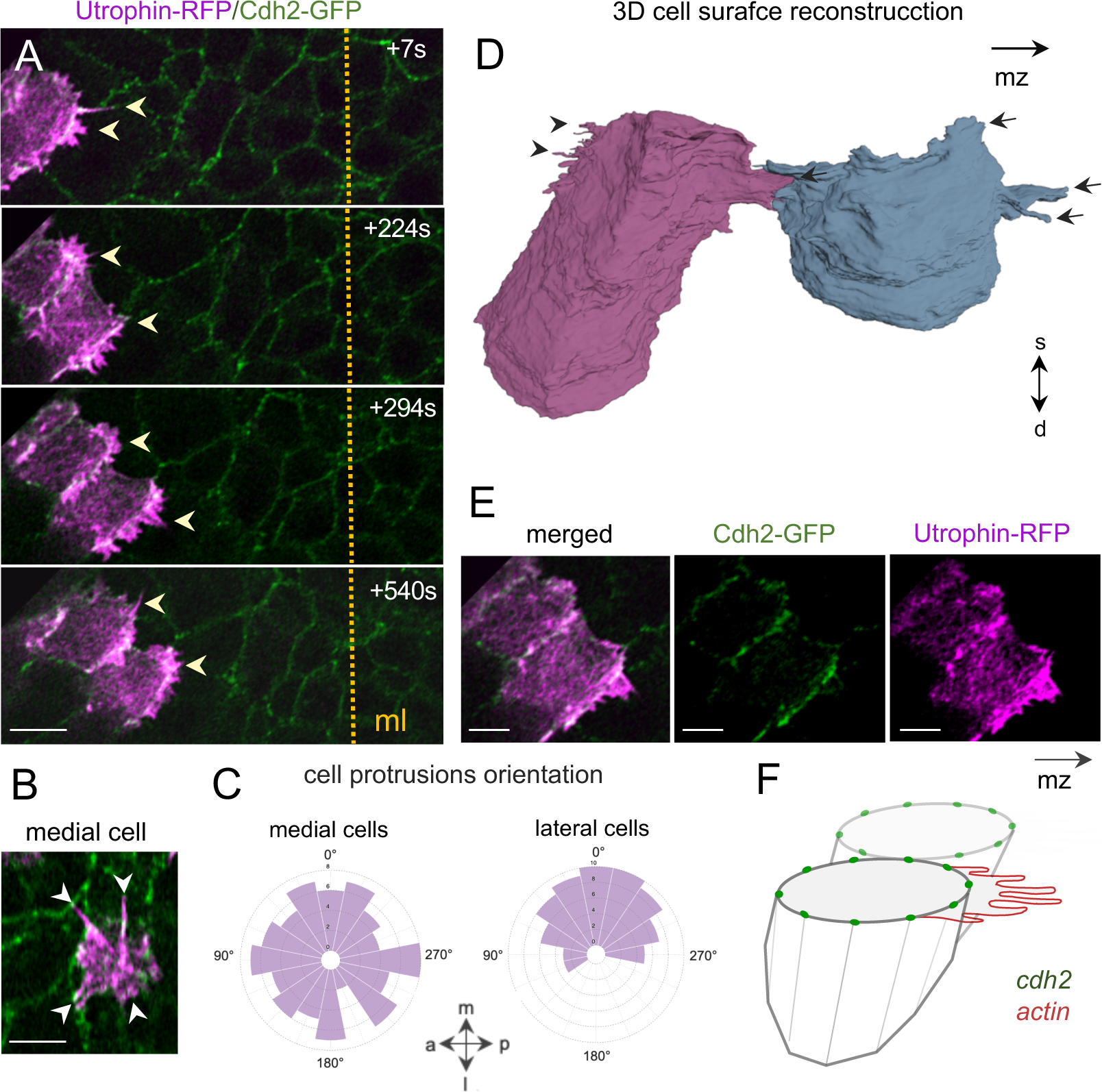
Superficial actin-rich protrusions. (A) Four frames from time-lapse sequence showing two neural plate cells labelled with the actin-binding protein utr-RFP (magenta) in a Cdh2-GFP background at 11.5 hpf. Anterior is up, medial is right. Dotted line denotes embryo midline (ml) and arrowheads indicate medial-oriented actin-rich protrusions. Time in seconds (s), scale bar 10µm. (B) Cell in midline zone with more evenly distributed protrusions. Scale bar 10µm. (C) Rose plot showing orientation of cell protrusions in medial and lateral neural plate cells (bin size, 22.5, n= 25 cells from 3 embryos). (D) SBFSEM 3D reconstruction of two adjacent neural plate cells during convergence. Arrows indicate midline-oriented cell protrusions while arrowhead depicts posterior oriented cell protrusions. mz midline zone, s-d denotes superficial-deep axis. (E) Relative location of Cdh2-GFP and Utr-RFP show actin-rich protrusions extend beyond Cdh2 puncta. Scale bar is 5µm. (F) Schematic of actin-rich protrusions and Cdh2 puncta. mz indicates midline zone.

The polarity of actin protrusions changes when cells reach the midline zone (we define this as within 20µm of the absolute midline). Within the midline zone protrusions are more evenly distributed around the cells’ superficial perimeters (Fig. 3B, C).

### Oscillatory contractions at the neural plate surface

We have visualised the detailed behaviours of individual cells at the neural plate surface by imaging the superficial cell perimeters from the dorsal surface (Fig. 4A), either by imaging the Cdh2 puncta that sit at cell-cell interfaces or by imaging cell membranes directly using membrane-GFP. Using a frame interval of between 5 and 10 seconds, cell perimeters at the superficial surface were seen to move in an oscillatory fashion (Fig. 4B and Movie 2). Thus cells are pulling on their neighbours in a pulsatile manner. How this behaviour is coordinated across individual cells and between neighbouring cells is complex and we have identified several different behaviours among and between cells.

**Figure 4.**
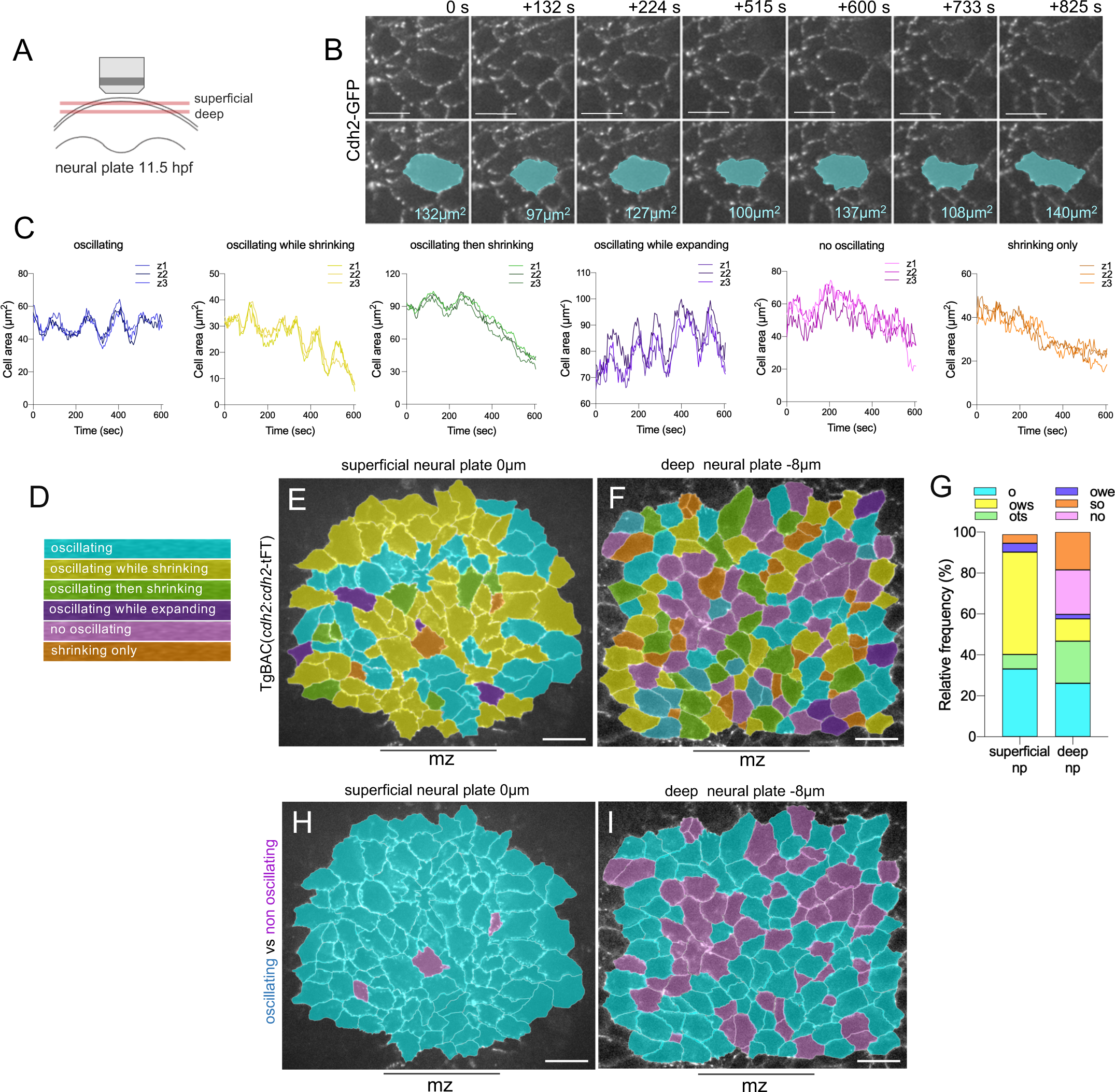
Oscillatory contractions. (A) Schematic of imaging set-up. (B) Selected time-lapse frames from superficial surface of neural plate to show dynamic cell profile. Bottom panels show changes in superficial surface area of selected cell (blue). Scale bar 10µm. (C) Examples of contractile oscillatory behaviours in superficial cell surface area. In each graph area is measured at three consecutive z-levels (0.5µm apart). (D) Colour code of distinctive cell surface behaviours. (E) Map of cell behaviours at superficial surface. (F) At deeper levels of the neural plate (z= 8µm from the surface), oscillatory contractile behaviour is less dominant and many cells show a non-oscillating dynamic. In E and F, anterior is up, mz indicates midline zone. (G) Histogram of cell surface behaviour at both superficial and deep levels of the neural plate. (H, I) Maps coloured to emphasize oscillating (blue) versus non-oscillating (purple) cells. mz is midline zone and scale bar is 20µm in E, F, H and I.

To understand these behaviours we analysed the superficial surface area of cells using either Cdh2-GFP or membrane-GFP labelled cells during an approximate 10 minute period. We identified and mapped six cell behaviours (Fig. 4C and Fig. S1). Cells that exhibit clear oscillations (repeated expansions and contractions of their superficial surface area) without an overall shrinkage of that cell surface. Cells that oscillate and also exhibit an overall shrinkage in surface area. Cells that oscillate without shrinking but then stop oscillating and shrink without oscillations. Cells that oscillate but also have an overall expansion of their surface area. Cells that show no clear oscillations but their surface area is noisy and may stay roughly the same size or shrink. The predominant behaviours of cells at the superficial surface are either oscillating (30%) or oscillating while shrinking (50%) (Fig. 4H). The period of oscillations for most cells was between 2 and 3 minutes and the amplitude was approximately 5 to 10% of a cell’s superficial surface area (Fig. S1). Oscillating cells were found in all regions of the hindbrain neural plate – i.e. both regions where cells converge to the midline and within the midline zone where cells internalise.

To determine whether the oscillatory behaviour was particular to the superficial surface of neural plate cells we repeated our analysis at a depth of 8µm below the superficial surface (Movie 6). In contrast to the superficial surface where only 5% of cell profiles showed no oscillatory behaviour, approximately 40% of cell perimeters analysed at 8 µm depth showed no oscillatory behaviour (Fig. 4F, G, I). Half of these non-oscillating perimeters showed shrinking without oscillations. Approximately 20 % of the deep cell perimeters show neither clear oscillations nor shrinking or expansion (Fig. 4F, G). For this 20% of cells their perimeters were not static but their movements were noisy, rapid and not easily defined. For the 60% of deeper cell profiles that did oscillate they showed similar oscillatory dynamics (frequency and amplitude) to those at the surface (Fig. S1).

### Myosin dynamics at the superficial neural plate surface

To help understand the contractile behaviours of cells at the superficial neural plate surface we analysed Myosin II activity and its distribution at the subcellular level using the Tg(*actb1*:*myl12. 1*-GFP) line. Myosin II fluorescence was very dynamic and largely appeared in three states. Cells had a uniform distribution of low intensity cytoplasmic fluorescence in addition to high intensity fluorescence organised in either a dynamic fibrillary manner or in high intensity dynamic foci with a stellate appearance (Fig. 5A and Movie 7). In the cells outside the midline zone, the fibrillary Myosin II was predominantly arranged in a mediolateral orientation, usually at the cell perimeters (Fig. 5B). For cells within the midline zone the fibrillary Myosin II was more randomly arranged (Fig. 5B). High intensity Myosin II foci were seen throughout the superficial surface, most often at cell perimeters and often at multicell vertices. Myosin II foci with radiating fibrillary Myosin II were also seen within the centre of the superficial surface of cells. Fibrillary myosin was longer within the paramidline zone and myosin foci were more frequent in the midline zone (Fig. 5C, D). The number of visible Myosin II foci was more dynamic in the midline zone than the paramidline zone and the duration of Myosin II foci was longer in the midline zone (Fig. 5E, F).

**Figure 5.**
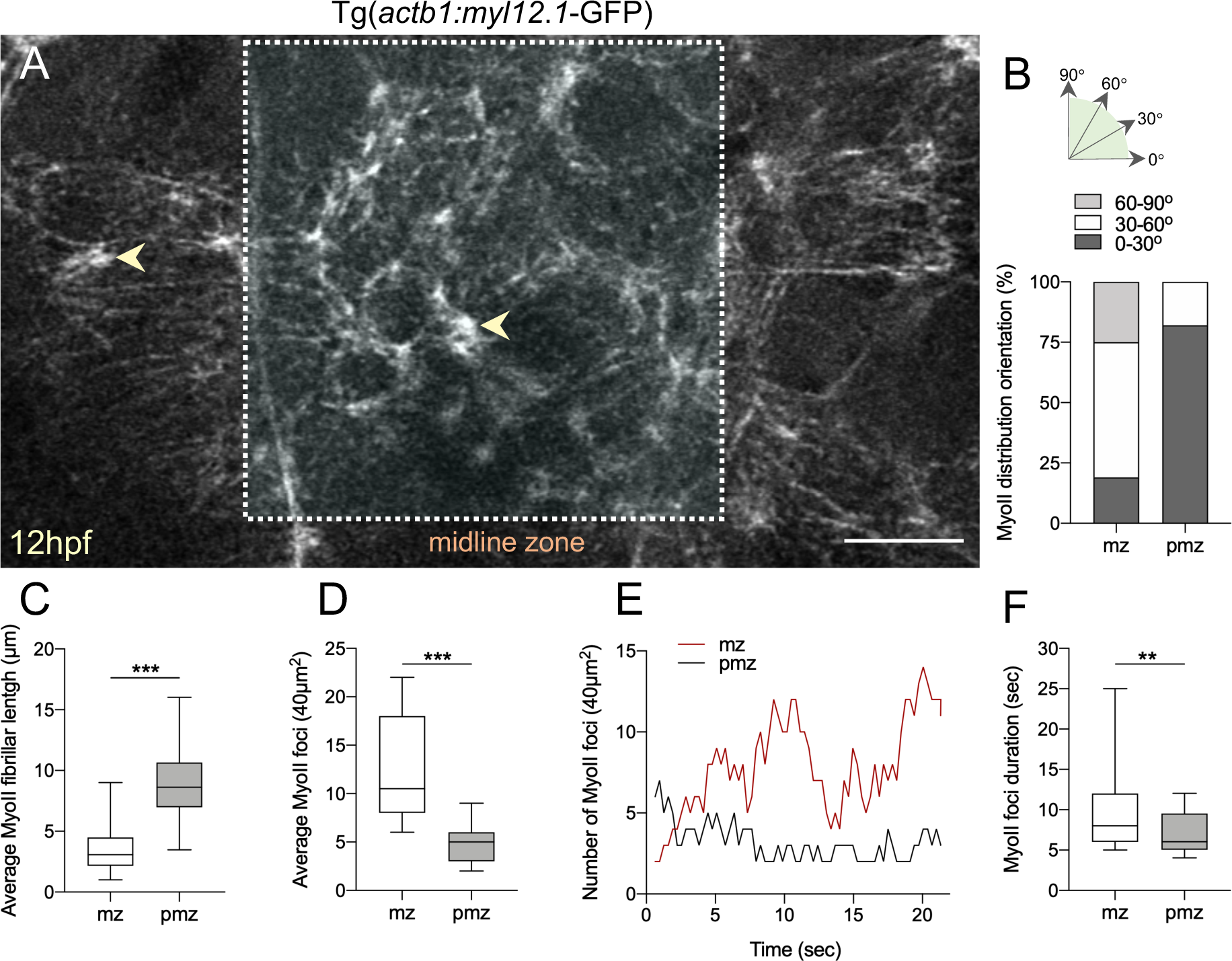
Myosin II activity at the superficial surface of the neural plate. (A) Maximal intensity confocal projection from a Tg(actb1:myl12.1-GFP) embryo at 12hpf. Arrowheads indicate puncta. Anterior to top, scale bar 10µm. (B) Fibrillar MyoII orientation in midline zone (mz) and para-midline zone (pmz). Two embryos analysed. (C) Length of fibrillar MyoII in midline zone (mz) and para-midline zone (pmz). 75 fibrillar structures analysed in each zone; ****P<0.0001. (D) Number of MyoII foci (stellate organization) in midline zone (mz) and para-midline zone (pmz). 25 foci analysed in each zone; ****P<0.0001. (E) MyoII foci number in midline zone (mz, red line) and para-midline zone (pmz, black line) over 20 mins. (F) Duration of MyoII foci in midline zone (mz) and para-midline zone (pmz). 15 foci tracked in each zone; ****P<0.0001.

### Neural plate dynamics and organisation in cdh2***^fr7/fr7^***mutants

Loss of Cdh2 function results in a strong neural plate phenotype as previously reported. Convergence of the neural plate to the midline is largely inhibited and very few midline cells internalise (Lele et al., 2002; Hong and Brewster, 2006; Biswas et al., 2010; Edmond et al., 2011; Araya et al., 2019). But how loss of Cdh2 function results in this phenotype is not clear.

Neural plate cells undergo two distinct behaviours in order to transition from plate to neural keel. They must converge to the midline and they must shrink their superficial surface area in order to internalise. During our 10 minute observation periods, 96% of all wildtype (wt) cells oscillate and 82% of cells that shrink do so while oscillating. This could mean oscillatory contractions are an important driver of both convergence and shrinkage. To begin to understand causality in these relationships we quantified the oscillatory behaviours of cells in *cdh2*^fr7/fr7^ mutants (Fig. 6A-H and Fig. S2A, B, C). We find that 92% of all neural plate cells in mutants still oscillate (Fig. 6D, E, F) and do so with the same periodicity and amplitude as wt cells (Fig. 6G, H). However, in mutants a smaller percentage of cells shrink (64% in wt compared to 45% in cdh2 mutants) and cells in the midline zone remain large compared to wt (Fig. S2D, E). The percentage of cells that shrink while oscillating decreases from 40% in wt down to 20% in mutants, while the percentage of cells that shrink without oscillating is essentially the same (24% in wt and 25% in mutants). Additionally, more oscillating cells expand in mutants (10% in wt compared to 19% in mutants). These data show that when Cdh2 adhesion is defective convergence is inhibited despite continued oscillatory contractions, and that oscillating cells are less able to shrink their superficial surface if they are Cdh2 defective.

**Figure 6.**
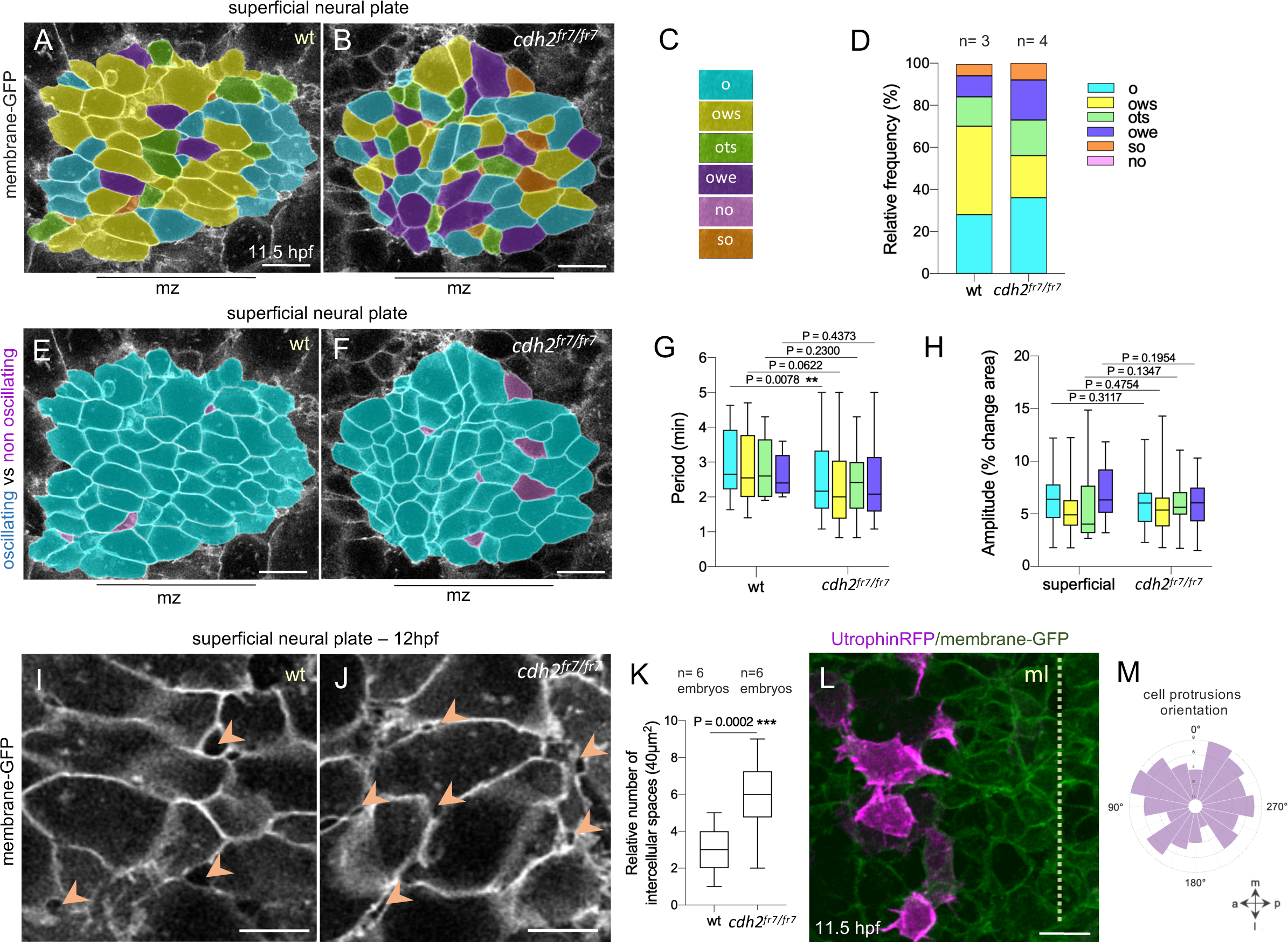
Neural plate surface dynamics in cdh2 muta. (**n**A**t**,**s** B.) Maps of oscillatory activity in wt and cdh2 mutants at 11.5 hpf. Scale bar 10µm. (C) Colour code for (A) and (B); o (oscillating), ows (oscillating while shrinking), ots (oscillating then shrinking), owe (oscillating while expanding), no (not oscillating), and so (shrinking only). (D) Frequency of cell surface behaviours in wt (3 embryos) and cdh2 mutants (4 embryos). (E, F) Colour code map showing distribution of oscillating (pale blue) vs non-oscillatory (pale purple) cells in wt and cdh2 mutants. mz indicates midline zone and scale bar is 10µm. (G) Box plot analysis comparing oscillation period between wt and cdh2 mutants. Three wt and four cdh2 embryos were analysed, bars indicate max and min values. (H) Box plot analysis comparing amplitude of oscillations in wt and cdh2 mutants. (I, J) Intercellular spaces (arrowheads) in wt and cdh2 mutants at 12hpf. Scale bar 5µm. (K) Box plots comparing number of intercellular spaces in wt and cdh2 mutants within a 40µm^2^ area, ****P<0.0001. (L) Maximum projection of utr-RFP labelled cells (magenta) in a cdh2 mutant. Dotted line shows midline. Scale bar 10 µm. (M) Rose plot analysis depicting orientation of cell protrusions in converging cdh2 mutant cells (bin size, 22.5°, n= 25 cells from 3 embryos).

Analysis of Cdh2 mutants revealed two other changes at the superficial surface of the neural plate. One is an increase in extracellular space (gaps) between the superficial cells (Fig. 6J, K, L, and Movie 8) and the second is that the actin rich protrusions are less polarised towards the midline than in wt cells (Fig. 6M, N, and Movie 9).

Together these results show Cdh2 adhesivity is not required for oscillatory contractions in neural plate cells but is a significant component of the mechanism that allows oscillating cells to converge and to shrink. In addition, Cdh2 adhesivity is required for the compactness of neural plate cells and is required for the medially biased polarity of actin-rich protrusions at the surface of the neural plate.

### Relationship between neural plate and the overlying EVL

Light microscope images and time-lapse movies show that the superficial surface of the neural plate is very close to and possibly in contact with the deep surface of the embryo’s protective epithelium - the enveloping layer (EVL) (Movie 10). This raises the question of whether the rapidly moving neural plate adheres to the stationary EVL and what cell behaviours might allow this dynamic interaction. To examine this relationship we used high resolution serial block face scanning electron microscopy (SBFSEM) imaging (Fig. 7). This reveals that, in regions outside the midline zone, the deep surface of the EVL and the superficial surface of the neural plate are in intimate, near continuous contact with very little intervening extracellular space and no obvious extracellular matrix (Fig. 7A-D and Movie 11). In contrast, at early neural keel stages, while no extracellular space is obvious between EVL and neural plate in paramidline locations, a distinct increase in extracellular space between the neural plate and the EVL has developed at the midline zone where cells are internalising to form the neural keel (Fig. 7E, F).

**Figure 7.**
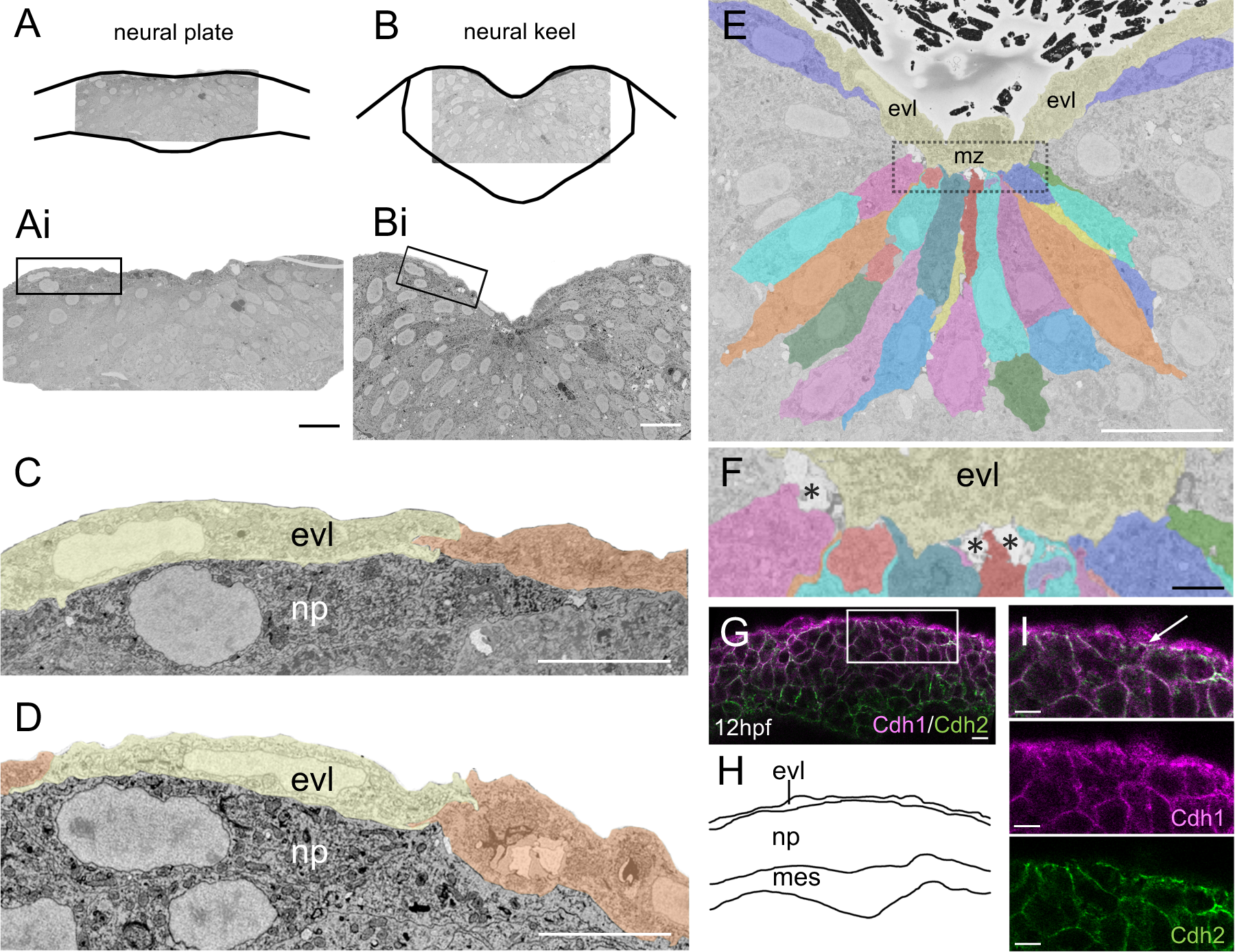
Relationship of neural plate and EVL. (A, B) Schematics to show tissue orientation in serial block face image Movies S12 and S13. (Ai, Bi) Low-mag micrographs showing regions of interest magnified in (C) and (D). Scale bar 20µm. (C, D) Close apposition of neural plate and EVL cells at plate and early neural keel stages. Scale bar 20µm. (E) Serial block face micrograph at 12hpf depicting relation of neural plate cells with EVL in the midline zone (mz, box). EVL cells are tinted yellow. Scale bar 20µm. (F) High mag of boxed region in (E). Asterisks indicate extracellular space. Scale bar 2.5µm. (G) Transverse confocal section of the neural plate at 12hpf stained for Cdh1 (magenta) and Cdh2 (green). Scale bar 10µm. (H) Schematic showing tissue organization in (S). (I) High mag of inset in (G), showing relative location of Cdh1 and Cdh2 at the EVL-neural plate interface.

To examine the possibility that neural plate and EVL cells adhere to one another we analysed the distribution of the Cdh1 adhesion molecule. In contrast to Cdh2 which is not expressed in EVL, Cdh1 immunoreactivity is found at the surface of both EVL and neural plate cells and enriched at their interface (Fig. 7G, H, I) suggesting homophylic (Cdh1-Cdh1) and heterophilic (Cdh1-Cdh2) adhesions could help stick these tissues together across this interface.

The close proximity and common adhesion molecule suggest EVL and neural plate are in intimate contact, but since neural plate cells “glide” past the stationary EVL cells (Movie 10) this interaction must be weak enough to allow rapid relative movement. To better understand how the stationary EVL cells behave in relation to the moving neural plate cells, we imaged the deep surface of the EVL to see if it would reveal any details of how these two tissues interact. Using a reporter of actin distribution in the EVL (Tg(Krt18:lifeact-RFP, Anton et al., 2018), we found the deep (basal) surface of the EVL is rich in dynamic, actin-rich lamellipodial protrusions (Fig. 8A, B and Movie 12). These EVL protrusions and their dynamics are also clearly visible (Fig. 8C, D and Movie 13) using the GT(ctnna-citrine)ct3a gene trap line (Zigman et al., 2011). This reporter shows Ctnna is abundant in the EVL protrusions (Fig. 8E), and enriched in the neural plate /EVL interface (Fig. 8C-F) and likely functions here to link Cadherins to the cell cytoskeleton. The Ctnna reporter does not appear to be expressed on the superficial lamellae or filopodial protrusions from neural plate cells but is expressed in the in puncta at the superficial junctions between neural plate cells and more generally in the cell membranes between neural plate cells. SBFSEM observations show the EVL protrusions lie in the plane of the interface between the EVL and neural plate (Fig. 8g and Movie 11).

**Figure 8.**
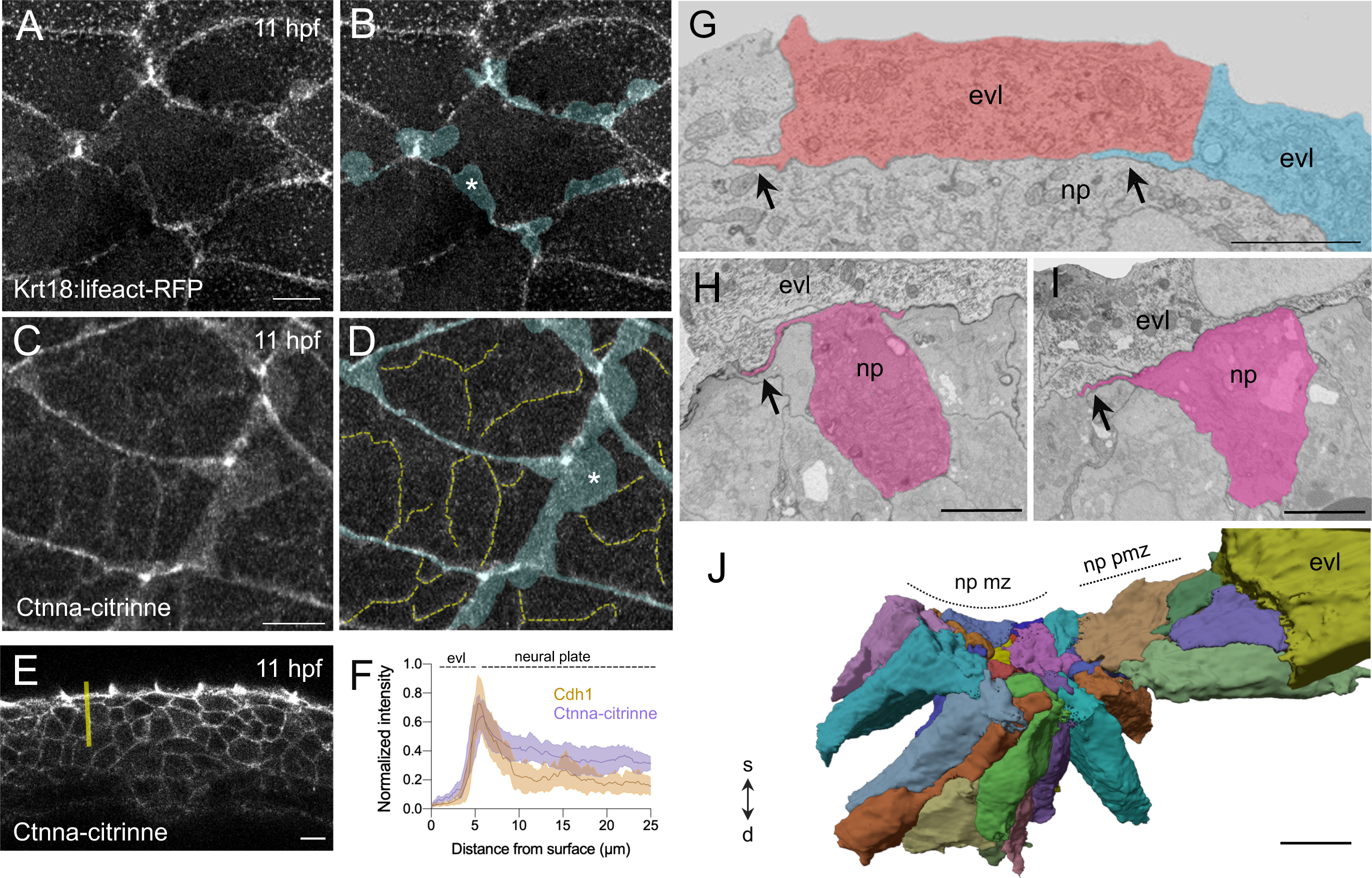
EVL and neural plate cell projections. (A, B) Single confocal image showing basal EVL lamellar projections (pseudo-coloured blue in B) in Krt18:lifeact-RFP embryos at 11hpf. Scale bar 10µm. (C, D) Maximum intensity projection at EVL-neural plate interface from a Tg(ctnna-citrine)^ct3a^ embryo at 11hpf. Scale bar 10µm. Dashed yellow lines in D denote Ctnna expression in neural plate cells and blue tint denotes Ctnna in EVL lamelli. Asterisks in B and D denote basal lamellas. (E) Transverse confocal section of the neural plate and EVL in a Tg(ctnna-citrine)^ct3a^ embryo at 11hpf. Yellow line indicates example of line selection for quantification of Ctnna intensity. Scale bar 10µm. (F) Normalised intensity of Cdh1 (brown) and Ctnna (purple) signal across EVL and neural plate. (G) Serial block face image across neural plate-EVL interface in para-medial region depicting medially-orientated basal projections of EVL cells (arrows). Scale bar 5µm. np is neural plate, medial to left. (H, I) Serial block face images across neural plate-EVL interface in para-medial region depicting medially oriented protrusions from neural plate cells (tinted pink). Medial to left, scale bar 5µm. (J) 3D reconstruction from serial block face images showing superficial surface of neural plate cells in midline (mz) and para midline (pm) zones. A single EVL cell is depicted in yellow, all other EVL cells have been removed from the reconstruction. S-D, indicates superficial-deep axis of the tissue. Scale bar 10µm.

Earlier we showed that the superficial surface of neural plate cells is characterised by medially biased lamellipodial protrusions (Fig. 3). SBFSEM imaging shows that, like the EVL protrusions, these neural plate protrusions also lie sandwiched along the plane of the EVL/neural plate interface (Fig. 8H, I and Movie 14). Three-dimensional reconstruction of multiple neural plate cells confirms their superficial surface is much reduced in area and more uneven at the midline zone compared to the larger and smoother surfaces of para-midline zone cells (Fig. 8J).

Combined with the observations of superficial protrusions from neural plate cells, these observations show the close apposition between neural plate and EVL cells is characterised by highly dynamic lamellipodial protrusions from both tissue layers. The protrusions from both layers lie sandwiched in the plane of the interface and do not protrude into the cells of the opposing layer. The distribution of Cdh1 suggests EVL and neural plate cells are likely to adhere to one another through homophylic adhesions and heterophilic Cdh1-Cdh2 adhesions across the tissue interface may also be possible (Ounkomol et al., 2010; Straub et al., 2011; Labernadie et al., 2017; Warga and Kane, 2018).

## Discussion

In this work we analyse the cell and subcellular behaviours that accompany movements of the zebrafish neural plate as it converges to the midline and internalises to form the neural keel. Our results show that during convergence of the neural plate to the midline cells move as a coherent sheet with little neighbour exchange. At the superficial surface cells are mechanically coupled together through Cdh2-rich punctate adhesions and converging cells almost continually tug on their neighbours through oscillatory surface contractions. Once in the midline zone the cells no longer behave as a coherent sheet but individual cells reduce their superficial surface area and internalise in a scattered fashion. Cells continue to oscillate in the midline zone and most internalising cells shrink their superficial surface area while oscillating, although a small percentage shrink without oscillations. Neural plate cells in cdh2 mutants are less compact and although they still oscillate they are unable to converge to the midline or reduce their superficial surface to internalise. Here we discuss the contributions that cell-cell rearrangements, oscillatory contractions, Cadherin based adhesions and polarised cell protrusions may have during this morphogenetic process.

### Zebrafish neural plate cells converge as a cohesive sheet

The motive forces that drive cells in the teleost neural plate to the midline are not well understood. In common with other vertebrates efficient narrowing of the neural plate requires signalling of the Planar Cell Polarity (PCP) pathway (Ciruna et al., 2006; Tawk et al., 2007) but how this controls cell and tissue movements is not certain. In other vertebrates and other tissues PCP function during tissue narrowing is often associated with cell intercalation along the mediolateral axis in a manner that simultaneously effects mediolateral narrowing and anteroposterior elongation (thus driving convergent extension). But here we show the collective narrowing of the zebrafish neural plate (at least during the phase of plate to keel transition in the hindbrain region) does not involve mediolateral cell intercalation, rather cells move as a cohesive sheet towards the midline with very little neighbour exchange. Here we have only analysed the prospective hindbrain region, and while mediolateral intercalation also contributes little to convergence in Xenopus anterior neural plate it does appear significant in more posterior regions (Christodoulou and Skourides 2022). When cells reach the midline zone, this cohesive sheet behaviour changes and cells now internalise as individuals in a scattered fashion (this work and Araya et al., 2019) to deepen the neural primordium at the midline and form the neural keel. Cell internalisation at the midline is itself a cell interdigitation event, but does not appear to be critically regulated by the PCP pathway since although PCP depletion decreases the speed of neural plate convergence, internalisation to initiate neural plate to neural tube transition still occurs (Tawk et al., 2007).

### Cdh2 adhesions

In addition to PCP signalling zebrafish neural plate morphogenesis is strongly dependent on the function of the adhesion protein Cdh2 (Lele et al., 2002; Hong and Brewster, 2006; Araya et al., 2019). Neural plate cells in Cdh2 loss of function mutants are unable to converge to the midline or internalise (Araya et al., 2019). Cdh2 mutant cells transplanted into a wild-type background accumulate dorsally in the neural tube (Lele et al., 2002), suggesting either they cannot move medially with their wild-type neighbours in the plate or they cannot internalise when they reach the midline zone (or both). Here we show Cdh2 protein has a particular dynamic punctate distribution around the superficial surface of all neural plate cells. We assume these puncta are composed of Cdh2 rich adhesions representing sites of strong homophylic adhesions between adjacent cells (McGill et al., 2009; Li et al., 2021) and mechanically couple these cells so they can converge to the midline as a cohesive sheet. This view is supported by our finding of increased extracellular space between Cdh2 deficient neural plate cells that suggests a loss of cohesion between neural plate cells. Increased space between cells has previously been reported in other Cdh2 deficient tissue (Mongera et al., 2018) and Cadherin functions to aid cohesion during *C.elegans* neurulation (Barnes et al., 2020). Since our current time-lapse analysis shows superficial NP cells almost continuously tug on each other in an oscillatory manner, we speculate a loss of adhesion between Cdh2 deficient cells in zebrafish results in a loss of the traction and the propagating motive forces required for convergence to the midline.

### Oscillatory behaviour

Oscillatory contractions appear to be a common feature of many morphogenetic processes across many animal groups (Martin, 2010; Gorfinkiel and Blanchard, 2011; Gorfinkiel, 2016; Miao and Blankenship, 2020) including Xenopus neural plate (Shindo et al., 2019). They have previously been briefly described in the anterior neural plate of the zebrafish (Werner et al., 2021) and here we quantify their dynamics and relate them to other behaviours at the surface of the zebrafish NP in the hindbrain region. The period of neural plate contractile oscillations is roughly 2.5 minutes and fits well with the oscillatory behaviours of other systems (Solon et al., 2009; Gorfinkiel and Blanchard, 2011). They are a more prominent feature of the superficial surface of neural plate cells than other parts of the cells deeper to the surface. They are prominent both in cells converging to the midline and in cells within the midline zone during internalisation. Although oscillations are a common feature of shrinking cells, they do not appear to be a required feature of the neural plate cells’ ability to shrink their superficial surface in preparation for internalisation at the midline, as shrinkage can occur without oscillations in a small percentage of cells. Thus the contractile mechanisms of oscillations and overall shrinkage can be independent of each other and able to work simultaneously or separately. Oscillations still occur with the same amplitude and frequency and in a similar proportion of cells in Cdh2 mutants, showing that Cdh2 mediated adhesion is not required for oscillation and suggests oscillations could be a cell autonomous feature of superficial neural plate cells.

While we currently do not understand the function of oscillatory contractions during convergence and internalisation, evidence in invertebrate systems suggest that cell oscillation is associated with the formation and transmission of biomechanical properties across the tissue (Martin et al., 2009; Solon et al., 2009). During *Drosophila* mesoderm invagination, asynchronous oscillations of apical actin-myosin is associated with the formation of supracellular actomyosin networks that confers the tensile properties across the tissue required for bending (Martin et al., 2010). We speculate that the oscillatory tugging between cells that adhere through Cadherin based adhesions are required to generate the tensile strength necessary for convergence movements of the neural plate towards the dorsal midline.

### Polarised actin-rich protrusions

Our analyses of cell morphology using both high resolution time-lapse imaging and serial block face reconstruction reveal that the surface each cell at the surface of the neural plate is morphologically polarised with dynamic lamellipodial and filopodial protrusions directed towards the midline. These protrusions are prominent at the superficial cell surface and lie in the plane of the interface between neural plate and the overlying EVL. These protrusions appear not to express either Cdh2 or Ctnna proteins, so probably don’t mediate Cadherin based adhesion and this perhaps suggest a more sensory/exploratory role (detecting or responding to a mediolateral positional gradient?). Whatever their role, their location along the superficial surface of the neural plate suggests this is key location for events critical for convergence to the midline. Once cells are within the midline zone the polarity of these actin-rich protrusions is lost – the protrusions are still present but here they are elaborated all around the superficial surface of cells. We speculate that here they may switch their role from potentially enabling convergence to enabling internalisation – outside the midline zone their polarity suggests they may detect a medial to lateral positional gradient, when at the midline this gradient would disappear and the even distribution of actin-rich protrusions could reflect their new position and trigger a new behaviour (internalisation). In Cdh2 mutants (where cells do not converge to the midline) the polarised distribution of non-midline protrusions is lost (Fig. 6M, N), the protrusions are still present but no longer biased to the midline. This could suggest Cdh2 is required for cells to know their position relative to the midline, although an alternative possibility is that loss of movement towards the midline somehow allows a redistribution of protrusions. We note that neural plate cell protrusions have been described previously in zebrafish (Hong and Brewster, 2006) however the protrusive activity described in that work is likely to reflect the overall dynamic spindle shape of cells rather than the actin-rich lamellae and filopodia we describe here at the superficial surface of the plate.

### Cdh2 and internalisation at the midline

Wild-type cells reduce their superficial surface area in the midline zone to internalise but Cdh2 deficient cells do not do that. A reduction in a cell’s superficial surface area is presumably a prerequisite for internalisation. The prominent appearance and dynamics of Myosin II foci in the midline zone leads us to suggest the reduction in surface area involves an active contractile process, but currently we do not know whether contractions are in the internalising cell or in its surrounding neighbours or both. Internalising cells potentially leave a hole in the neural plate surface. This hole could be sealed simply by the shrinking cell drawing its neighbours together through cell-cell adhesions, or alternatively surrounding cells could purse-string together to seal over the potential gap (and potentially squeezing the internalising cell below the surface). A cooperative mechanism between surrounding cells has been proposed to seal tissue holes resulting from cell internalisation during *C.elegans* gastrulation (Pohl et al., 2012), and internalisation by capping over with neighbouring cells contributes to invagination in salivary glands and teeth (Li et al., 2020). We find more prominent gaps between cdh2 mutant cells and fewer mutant cells are able to shrink their superficial surface compared to wt. While 42% of cells in wt shrink while oscillating this is reduced to only 20% of cells in cdh2 mutants. This suggests Cdh2 adhesivity is an important component of the shrinking mechanism in oscillating cells. Additionally, since more oscillating cells in cdh2 mutants are able to expand this potentially suggests cells with normal Cdh2 adhesions are better able to stabilise a reduced superficial area. Perhaps the increased extracellular space between mutant neural plate cells simply allows more cells to expand into that space or the reduced adhesion and increased gaps between cells eliminates the cooperative mechanisms of internalisation suggested above.

### EVL/neural plate interaction

Unlike the superficial surface of the neural plate in all non-teleost species, the superficial surface of the teleost neural plate is not a “free” surface. It is covered by a thin squamous epithelium called the enveloping layer (EVL). Rather little attention has previously been given to the possible interactions between neural plate and the overlying non-neural EVL. Here we show the two tissues are in intimate contact, apparently without intervening extracellular matrix, and Cdh1 and Ctnna distributions suggest neural plate and EVL adhere to each other. Our antibody labelling suggest Cdh1 is expressed in membranes of both EVL and plate cells, so EVL/neural plate adhesions could be mediated by homophylic Cdh1 interactions, however since heterophylic interactions between Cdh1 and Cdh2 have been shown in mammalian cells (Straub et al., 2011; Labernadie et al., 2017) and suggested in zebrafish embryos (Warga and Kane, 2018) there may also be heterophylic adhesions between EVL and neural plate. None the less, EVL/neural plate adhesions must be either relatively weak or rapidly dynamic to enable the motile plate to move against the static EVL. Our time-lapse imaging show this tissue interface is an extremely dynamic environment with both neural plate and EVL cells elaborating dynamic actin-rich protrusions along the tissue interface. One possibility is that extension and retraction of these dynamic protrusions minimise the time that any individual adhesions can be retained between the two layers, i.e. they potentially act as an anti-friction mechanism between the two surfaces. It also seems likely that the EVL confines the movement of the converging neural plate cells – restricting the directionality of movement to the plane of the neural plate. Without the EVL the contractile oscillations of the neural plate cells could potentially squeeze cells or parts of cells up and out of the superficial plane of the neural plate. But with the arched EVL lying over and in contact with the neural plate, the surfaces of plate cells are constrained to a flat surface, an arrangement potentially mechanically beneficial in moving the plate to the midline. Tensile properties of non-neural ectoderm has previously been suggested to aid the movements of neurulation in Xenopus embryos (Morita et al., 2012; Christodoulou and Skourides, 2022), but in that case the non-neural ectoderm lies alongside the neural plate in contrast to the EVL which sits directly above the zebrafish neural plate. Basal actin-rich protrusions from zebrafish EVL cells have previously been reported much earlier in development, where they are suggested to aid the rearrangements of deep cells during epiboly (Rutherford et al., 2019). Finally, when cells reach the midline zone they internalize, so any adhesions between EVL and neural plate within this zone must be lost and this is potentially reflected in the increase in extracellular space that we observe between plate and EVL at the midline (Fig. 7E, F).

## Materials and Methods

### Animals

Adult zebrafish (*Danio rerio*) were kept at standard conditions at 28.5°C water with a 14 hour /10 hour photoperiod light cycle (Westerfield, 2000) at both the King’s College London (KCL) and Universidad Austral de Chile (UACh) Fish Facilities. Wild type AB, TgBAC(*cdh2:Cdh2*-tFT) (Revenu et al., 2014), *cdh2*^fr7/fr7^*/pac* (Lele et al., 2002), Tg(*actb1:myl12*.*1*-GFP) (Maitre et al., 2012), Tg(*actb1:GFP-utrCH*) (Behrndt et al., 2012), and Tg(*ctnna-citrine*)^ct3a^ (Zigman et al., 2011), Tg(Krt18:lifeact-RFP) (Anton et al., 2018) zebrafish lines were used in this study. Embryos were raised in E3 medium and staged by morphology according to Kimmel et al. (1995). Developmental stages were given in terms of hours post fertilisation (hpf). All animal experiments were performed in compliance with the guidelines and approved by animal ethics committees of both KCL (Home Office Animals, Scientific Procedures, Act 1986, ASPA), and by UACh/DID animal committee.

### mRNA synthesis and microinjection

pCS2+ expression vectors carrying membrane-localized fluorescent CAAX-GFP (mGFP) or Utrophin-RFP (utr-RFP) were linearized using restriction enzymes (Promega) for two hours at 37°C and precipitated at -20°C overnight in 70% ethanol. mRNA *in vitro* transcription was performed using the SP6 mMessage mMachine kit (Ambion, AM1340), purified through a column (Roche), and final concentrations were measured using a Nanodrop 3300 spectrometer (Thermoscientific). Embryo microinjections were performed under a dissecting microscope using a glass slide and petri dish (Westerfield, 2000). Microinjections were performed using a glass micropipette with filament (Harvard Apparatus) coupled to a micromanipulator and attached to a Picospritzer® (General Valve Corporation). mRNA was injected at 80-100 pg per embryo. For ubiquitous localization, mRNA was injected at one cell stage. For mosaic injection, mRNA was injected into a single blastomere at 128 cell-stage embryo.

### Live confocal imaging

Embryos at neural plate stages were mounted in 1.5% low-melting-point agarose in a glass-bottomed imaging chamber and filled with embryo medium at 28.5°C and supplemented with Tricaine at 0.05% (Sigma). Embryos were orientated dorsal uppermost at the level of the hindbrain. Images acquired on a Zeiss LSM 880 Fast Airyscan microscope provided with heated stage at 28.5°C, and imaging with water-dipping × 20/1.0 NA objective lens. Time-lapse images were most frequently taken with a frame interval of 5-10 seconds and z-stack depth of approximately 5 to 10 µm and 0.3µm z-resolution. All raw Airyscan images were processed with Zeiss Airyscan processing software.

### Cell rearrangements during convergence

Cohorts of approximately 50 cells arranged in roughly 4 rows were followed during convergence to midline. Changes to cohort dimensions were analysed by measuring the maximum anteroposterior and maximum mediolateral dimensions at each time point. Total cohort area was measured by outlining the cohort of cells at each time point.

### Imaging processing and Oscillation analysis

Imaging processing were carried out using ImageJ software. To track cell movements over time, we used the Manual Tracking plugin using the cell centroid as reference. Projections of z-stacks are maximum projections unless otherwise indicated. To measure the duration of time individual Cdh2-puncta were expressed at the cell membrane, movies with 5 second time intervals were used. Cdh2-puncta (n=105) were randomly selected and tracked frame-by-frame, from the frame of first appearance to the frame the Cdh2-puncta were last visible in. The number of frames the Cdh2-puncta was visible in multiplied by the imaging time interval generated the length of time the Cdh2-puncta was visible at the cell membrane. To count the number of puncta at the cell boundary over time, we selected cells (n=8) with the whole membrane boundary clearly visible for at least 30 frames. The cell boundary of these cells was individually traced manually using a segmented line and saved as ROIs (regions of interest). The ROIs were then converted into coordinates using a FIJI plugin code. The individual cell coordinates generated, and the movie file were then run through a customized code. The Cdh2-GFP intensity values were standardized in each movie by 10 dividing every intensity value by the largest intensity recorded in the movie, generating Cdh2-GFP intensity values between 0 and 1. In a 0.25µm width region around the cell boundary, intensity of any Cdh2-GFP expression is measured in ‘steps’ of 0.1194µm increments along the cell perimeter. Regions of high intensity which met the relative Cdh2-GFP intensity parameters (minimum peak prominence: 0.1 and minimum intensity value: 0.5) set to differentiate Cdh2-puncta from non-clustered Cdh2 expression and background noise were then highlighted. The absolute value for the number of Cdh2-puncta present at the cell boundary for each frame was counted. To quantify the number of Cdh2-puncta with respect to the length of the membrane, the cell perimeter was measured for the same cells previously used to count the number of Cdh2-puncta. The cell perimeter was then divided by the number of Cdh2-puncta present in that frame, calculated in microns per puncta, and an average across all time points was made for each cell. Pixel intensity quantification for Fig. 8C was carried out as previously described in detail (Araya et al., 2019). To extract cell plasma membrane dynamics over time, we used freehand tool to manually segment cell perimeter at different z-levels. To reduce noise in these measurements, control measurements were established by using repeatedly segmentation (50 times) of one random frame in analyzed cells. We found that highest amount variability in the data was about 5 μm^2^ and thus it was considered as noise rather than oscillations.

### Statistical Analysis

All statistical tests were performed with Graphpad Prism 9.5. For significance between mean values, the non-parametric Mann-Whitney post-test were used. In all figures, statistical significance for p-values (≥0.05) is indicated as follows: * p<0.05, ** p<0.01, and *** p<0.001. Unless indicated, all the error bars shown in figures are SEM (standard error of the mean). Graphs in figures were generated using GraphPad Prism 9.5.

### Tissue preparation for Serial Block Face SEM

Embryos were embedded in agarose prior to immersion in fixative (1.5 % glutaraldehyde, 2 % paraformaldehyde, 10mM sucrose and 1mM calcium chloride in 0.1 M sodium cacodylate buffer) at 4 degrees overnight. After several rinses with 0.1M sodium cacodylate buffer containing 1mM calcium chloride and 10mM sucrose, the samples were further fixed in 2% osmium tetroxide, 1.5% potassium ferrocyanide and 1mM calcium chloride in 0.1M cacodylate buffer for 1 hour at 4°C. Tissue was then thoroughly rinsed in distilled water and incubated in 1% aqueous thiocarbohydrazide for 5 min. After further rinsing, the samples were treated with 2% aqueous osmium tetroxide for 30 min at room temperature, rinsed and en-bloc stained in 1% uranyl acetate overnight at 4°C. To enhance the contrast, the samples were incubated with Walton’s Lead solution for 30 min at 60°C, before proceeding to dehydration in acetone series and infiltration with Durcupan ACM resin (Sigma). After polymerisation for 48 h at 60°C, tissue blocks were mounted on aluminium pins using conductive glue (CircuitWorks Conductive Epoxy) and trimmed accordingly. Before imaging, samples were gold coated to increase electron conductivity. The specimens were then placed inside a JEOL field emission scanning electron microscope (JSM-7800F) equipped with a Gatan 3View system and OnPoint detector (Gatan). Section thickness was set at 50 nm (Z resolution). Samples were imaged at 2 kV under high vacuum, with an image size of 4180 x 10174 and a final pixel size of 18 nm.

### 3D reconstructions from SBFSEM data

Raw collections of 2D SBFSEM images were aligned, adjusted and manually segmented using the ImageJ plugin TrakEM2 software (Cardona et al., 2012). For 3D reconstruction, generated raw models in ImageJ were exported to Blender software v.279 (www.blender.org) with Neuromorph toolset for morphometric analysis (Jorstad et al., 2015).

## Acknowledgments

We thank the members of the Clarke lab for many discussions of this work, especially Rachel Moore for help with embryo production and the zebrafish facility at King’s College London for their help in maintaining zebrafish stocks. We thank John Fadul and Jody Rosenblatt for the aliquot of the Cdh1 antibody.

---

**Supplementary Figure 1.**
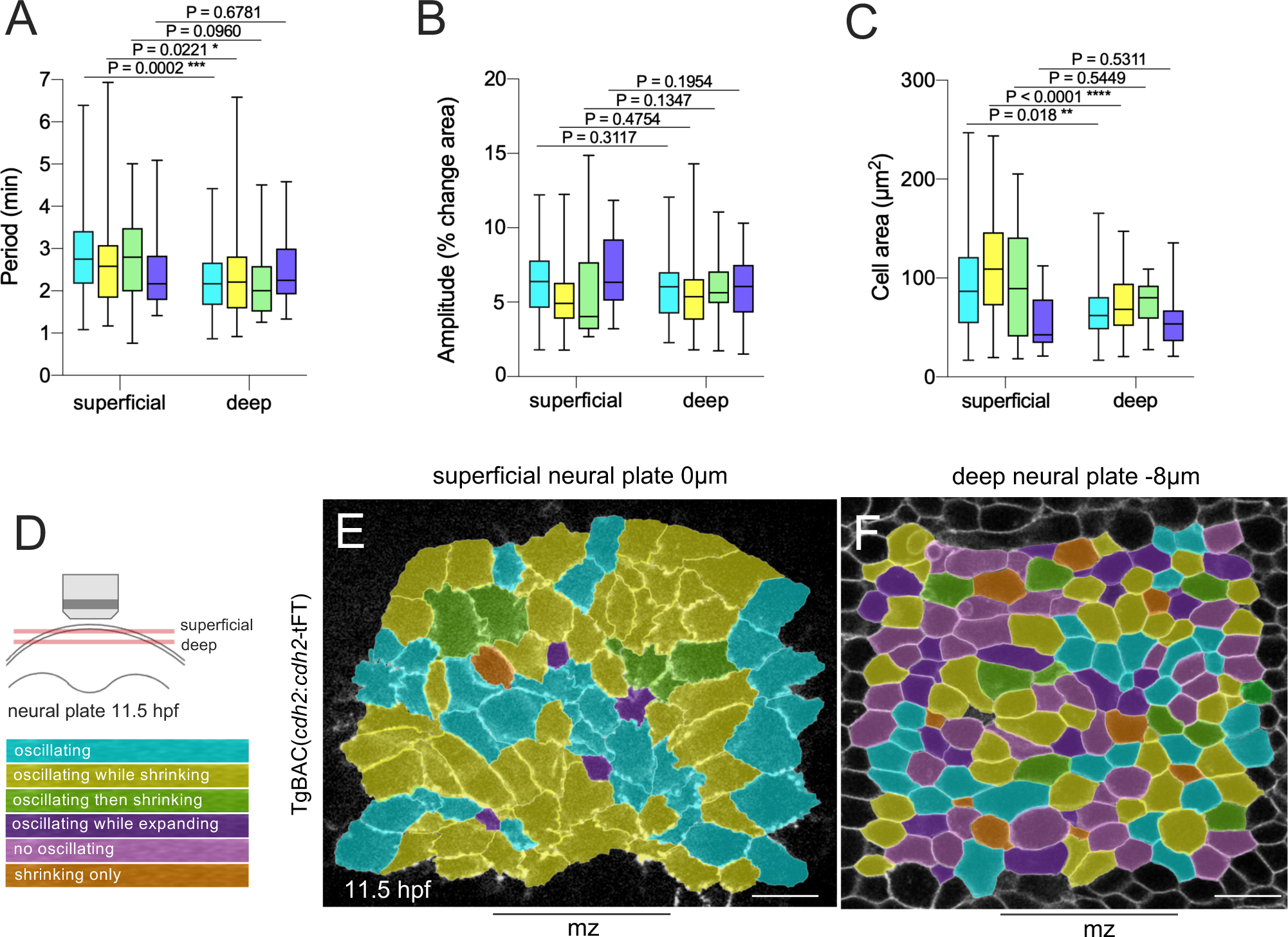

**Supplementary Figure 2.**
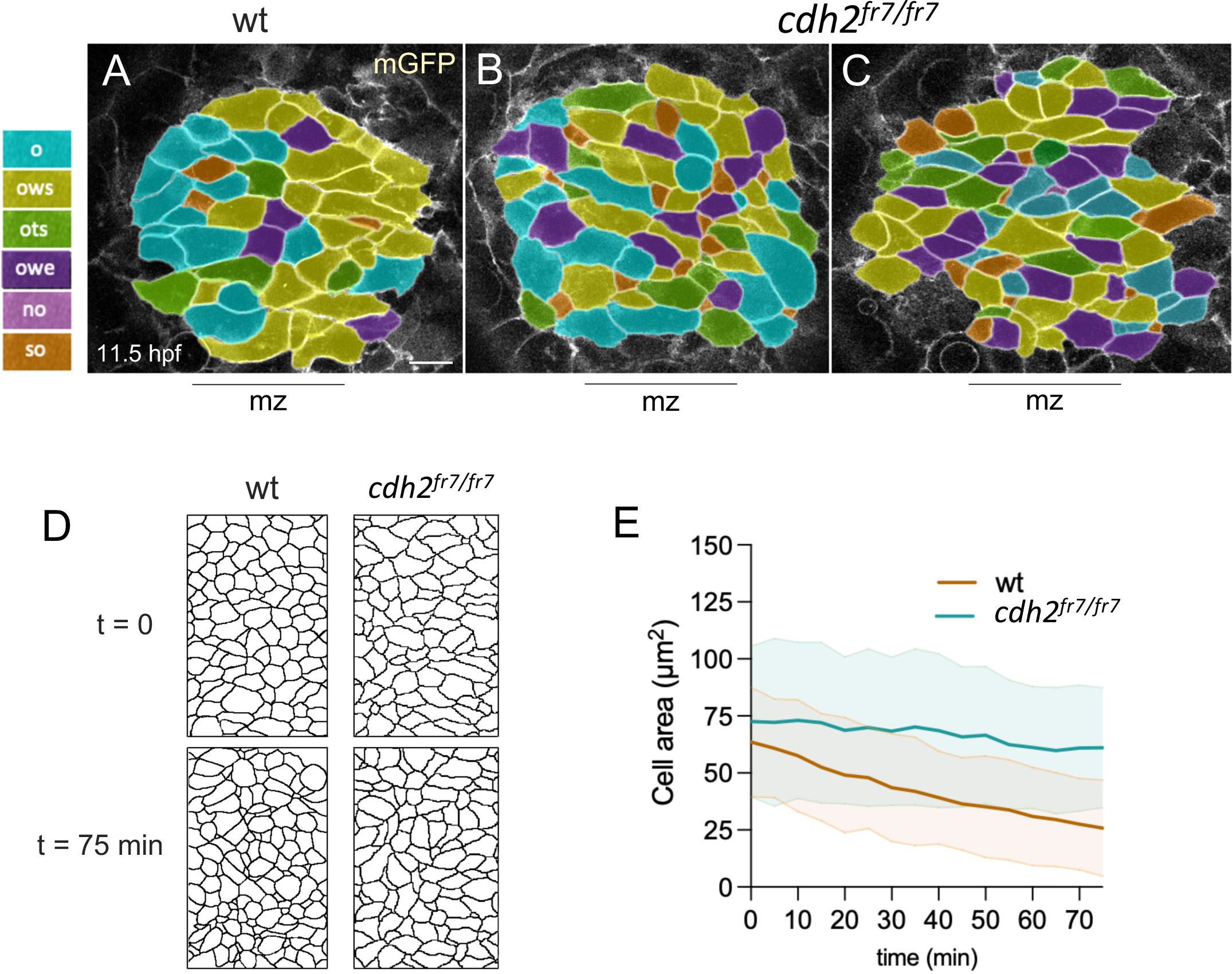

